# NanoARG: A web service for identification of antimicrobial resistance elements from nanopore-derived environmental metagenomes

**DOI:** 10.1101/483248

**Authors:** G. A. Arango-Argoty, D. Dai, A. Pruden, P. Vikesland, L. S. Heath, L. Zhang

**Affiliations:** Department of Computer Science, Virginia Tech, Blacksburg, VA, USA; Department of Civil and Environmental Engineering, Virginia Tech, Blacksburg, VA, USA

## Abstract

Direct selection pressures imposed by antibiotics, indirect pressures by co-selective agents, and horizontal gene transfer are fundamental drivers of the evolution and spread of antibiotic resistance. Therefore, effective environmental monitoring tools should ideally capture not only antibiotic resistance genes (ARGs), but also mobile genetic elements (MGEs) and indicators of co-selective forces, such as metal resistance genes (MRGs). Further, a major challenge towards characterizing potential human risk is the ability to identify bacterial host organisms, especially human pathogens. Historically, short reads yielded by next-generation sequencing technology has hampered confidence in assemblies for achieving these purposes. Here we introduce NanoARG, an online computational resource that takes advantage of long reads produced by MinION nanopore sequencing. Specifically, long nanopore reads enable identification of ARGs in the context of relevant neighboring genes, providing relevant insight into mobility, co-selection, and pathogenicity. NanoARG allows users to upload sequence data online and provides various means to analyze and visualize the data, including quantitative and simultaneous profiling of ARG, MRG, MGE, and pathogens. NanoARG is publicly available and freely accessible at http://bench.cs.vt.edu/nanoARG.

## INTRODUCTION

Antimicrobial resistance (AMR) compromises our ability to prevent and treat infectious disease, and presently represents one of the most significant and growing global public health threats (1). It is estimated that the annual number of deaths due to antibiotic resistance will top ten million by 2050 (2). In response, numerous national and international agents have called for increased monitoring, with increasing awareness of the importance of environmental monitoring. In particular, environmental monitoring can provide insight into not only human and agricultural inputs of antibiotic resistant bacteria and antibiotic resistance genes (ARGs), but also factors contributing to the evolution and spread of resistant pathogens. For instance, various environmental compartments, such as wastewater treatment plants, livestock lagoons, and amended soils can essentially act as “environmental reactors,” in which resistant bacteria discharged from domestic, hospital, industrial, and agricultural waste-streams have the opportunity to interact with native aquatic and soil bacteria in the presence of various selection pressure and give rise to new resistant forms (3,4). Humans may subsequently be exposed via consumption of food-crops affected via biological soil amendment or irrigation, as well as through contact with treated and untreated water used for recreational, hygienic, and potable purposes (5,6).

Molecular-based monitoring presents great advantage over culture-based techniques for tracking antibiotic resistance in the environment, particularly with respect to the potential to recover rich information regarding the carriage and movement of ARGs with complex microbial communities. Culture-based techniques, on the other hand, are highly time consuming and only provide information about one target species at a time, thus overlooking potentially key microbial ecological processes contributing to the spread of antibiotic resistance. Thus, directly targeting ARGs as “contaminants” of concern that transcend bacterial hosts has gained popularity. In particular, horizontal gene transfer plays a critical role in the rise of new resistant strains and dissemination of multi-antibiotic resistance. Correspondingly, multi-drug resistance has emerged as a major clinical challenge. For example, methicillin resistant *Staphylococcus aureus* (MRSA) is responsible for major hospital infections, with few options for treatment, especially when resistant to vancomycin (7). More recently, New Delhi Metallo beta lactamase (*bla*NDM-1) has emerged as major concern, encoding resistance to powerful last-resort carbapenem antibiotics and was carried on a highly mobile genetic element and associated with multi-drug resistance (8). Such examples emphasize that ideally, environmental monitoring technologies should provide a rapid and robust characterization of ARGs and their likely associations with mobile genetic elements, multi-drug resistance, and carriage by pathogen hosts. In this regard, shotgun metagenomic sequencing techniques have emerged as a promising tool for tapping into the diverse array of ARGs characterizing different environments (4,9-11). In particular, high-throughput next-generation DNA sequencing technologies, such as Illumina (12) and 454 pyrosequencing (13,14), have enabled a new dimension to ARG monitoring in the environment. While providing unprecedented amount of sequence information, a major drawback of these technologies is the very short DNA sequence reads produced, at most a few hundred nucleotides long. Still, next-generation DNA sequencing is growing in use as a powerful means of profiling ARG occurrence in various environments using either a direct annotation technique where sequences are compared against available ARG databases and relative abundances of the ARGs are then estimated, or an assembly based annotation technique where the short reads are assembled into longer contigs for ARG and also neighboring gene identification. Both approaches have limitations. The first is only applicable to previously-described ARGs populating available databases (15) and requires determination of an arbitrary DNA sequence identity cutoff (16), which undermines the possibility of identifying novel ARGs although a novel similarity based method has been proposed recently to annotate ARGs with low similarity to existing database ARGs (17). Assembly, on the other hand, requires deeper and more costly sequencing along with high computational resources (18) and still can produce incorrect contigs and chimeric assemblies (19). Thus, it is important to be cautious in interpreting results derived from assembly of short sequence reads because of possibility of assembly errors and lack of standard means to estimate confidence in accuracy of assembly (20-22).

In 2014, Oxford Nanopore Technologies (ONT) released the MinION nanopore sequencer that provides very long sequence reads averaging 5kb in length (23). It has been reported that nanopore can produce sequences with lengths even larger than 100kb (24). A major disadvantage of nanopore technology is the high error rate, estimated by Jain et al (2016) below 8% (25). However, this error rate represents a marked improvement over an earlier estimated error rate of 38% (26), with a general trend towards improved error rates with the help of read correction algorithms (27). A recent study demonstrated the potential for nanopore technology to produce highly accurate assemblies, with an accuracy of approximately 95% when applied to whole-genome sequencing (28-30). Nanopore sequencing has also been applied for the purposes of shotgun metagenomics, such as identification of viral pathogens (31), assessment of microbial diversity in extreme environments (32), and detection of ARGs (33-38). To date, nanopore sequencing has not been applied for the purpose of metagenomic profiling of ARGs in environmental samples.

Here we introduce NanoARG, a user-friendly online platform that enables comprehensive metagenomic profiling of a diverse range of ARGs and other relevant genes in environmental samples using nanopore sequencing. In addition to comprehensive ARG profiling, NanoARG also provides annotation of metal resistance genes (MRGs), mobile genetic elements (MGEs), taxonomic markers, and pathogens, along with interactive visualization of linkages among these various elements on the same DNA sequences. To demonstrate the potential of NanoARG for environmental ARG profiling, several MinION nanopore sequencing libraries, including environmental and clinical samples, were analyzed. The Web service is freely available at http://bench.cs.vt.edu/nanoARG/. It requires a user login and subscription to upload and process nanopore sequencing data.

## METHODS

### Web Service and Pipeline

**Figure 1** illustrates the NanoARG architecture. The workflow has three major components: 1) a Web interface, where users can upload data and monitor the progress of the analysis (**Figure 1A**); 2) a RESTful (REpresentational State Transfer) application program interface (API), which monitors and sends the raw MinION nanopore sequencing data to a computing cluster for processing (**Figure 1B**) and retrieval of results and downstream analyses, such as taxonomic annotation, gene co-occurrence analysis, human pathogens detection, network analysis, and multiple sample comparisons (**Figure 1C**). Presently the nanopore reads are screened against different databases using different omics tools, but this could be adjusted in the future as databases and annotation tools improve. Results are stored as Javascript Object Notation (JSON) files. Metadata and user information are encrypted and stored in a Mongo database. In addition, the entire workflow runs on a large distributed system in the Advanced Research Computing (ARC) center at Virginia Tech. The cluster is managed by the qsub queuing system (39).

**Figure 1:**
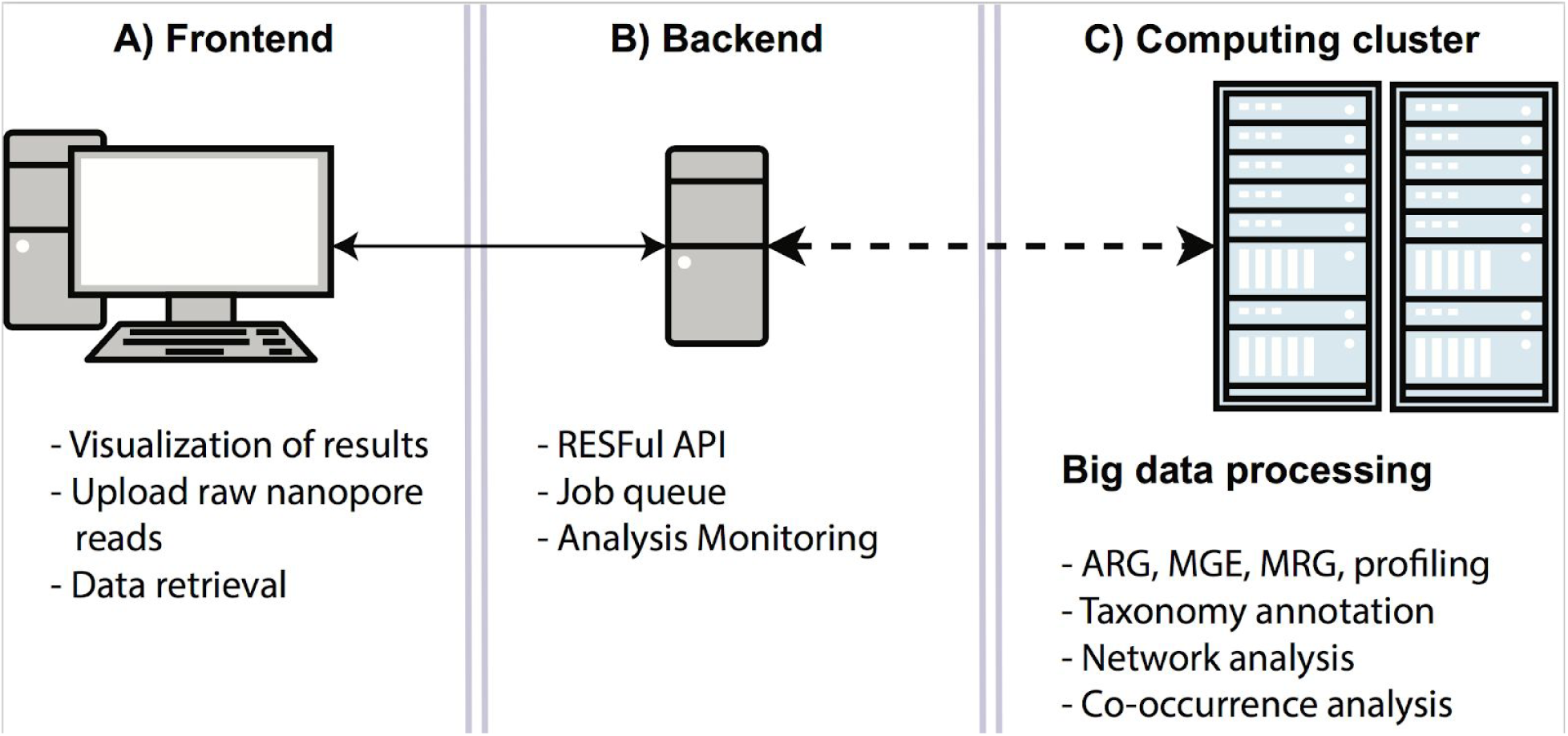
the NanoARG architecture. **A)** Frontend is the link between the users and the analysis by allowing users to upload raw data and visualize results. **B)** A backend RESTful API that manages the data, triggers the analysis and monitors the status of the analysis. **C)** Computing cluster is the module that process the data with ARG, MGE, MRG and taxonomy profiling.

The Web service provided by NanoARG includes several features to facilitate the analysis of environmentally-derived metagenomic data obtained from nanopore MinION sequencing. Users can submit data to the NanoARG Web service using a simple graphical user interface (**Figure 2A**). In the current version of NanoARG, data submitted to the system is stored privately. To start using the service, users are required to register an account, which allows users to manage and control submitted samples and projects. Users can voluntarily share their projects with other users by simply adding their email addresses. To create a project, a few parameters, such as name, description, and biome (**Figure 2B**), are required. Inside each project, users can add new samples, run new analyses, or remove or rerun old samples (**Figure 2C**).

**Figure 2:**
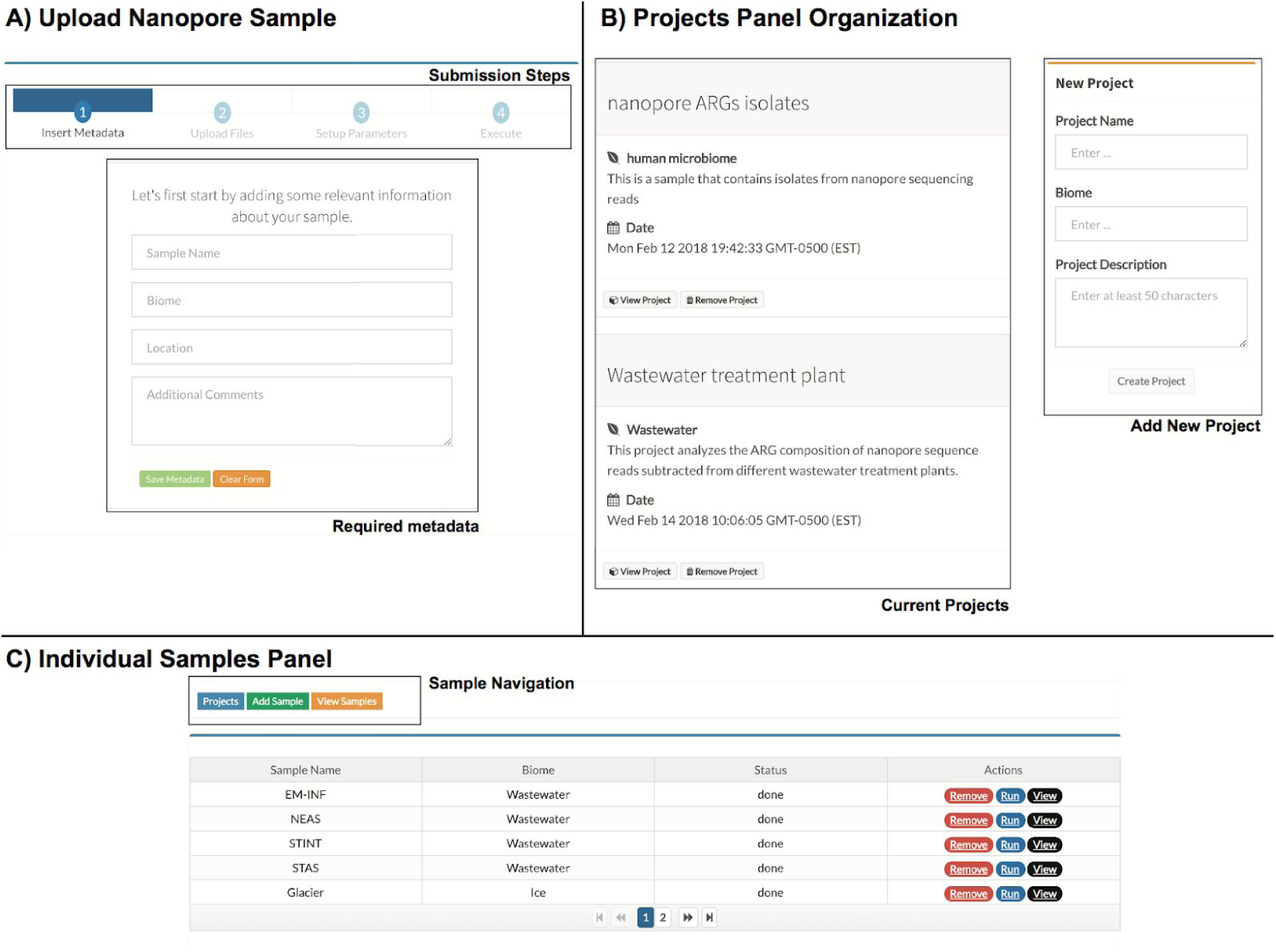
User Interface. **A)** steps and metadata required to upload samples to NanoARG. **B)** Projects are organized based on the creation date and visualized as a timeline post. **C)** List of samples under a project displaying basic metadata (Biome), the monitor variable (Status) and the three actions that can be performed by users.

### Required Data Types

NanoARG requires users to upload nanopore reads in FASTA format (40), assuming that the users have already preprocessed the raw fast5 files from the MinION device. This step can be done by using a base-calling program such as Albacore (41), Metrichor (23), or Nanocall (42), with a sequence extractor toolkit such as porertools (43). Barcode recognition and read sorting by barcodes can be conducted along with base calling. Before submitting data to the system, users are required to provide simple metadata consisting of sample name, biome, location, and comments and can also manually enter details about the sample extraction if necessary. Then following four simple steps (insert metadata, upload files, set up parameters, and execute), users can submit raw data and initiate the analysis (**Figure 2A**).

### Data Processing

Once raw data is uploaded to the computing cluster, it is processed by several modules that perform a set of tasks to obtain annotation profiles for ARGs, MGEs, MRGs, and associated taxa (**Figure 3**). Status of the analysis can be easily monitored through the user interface (**Figure 2C**).

**Figure 3:**
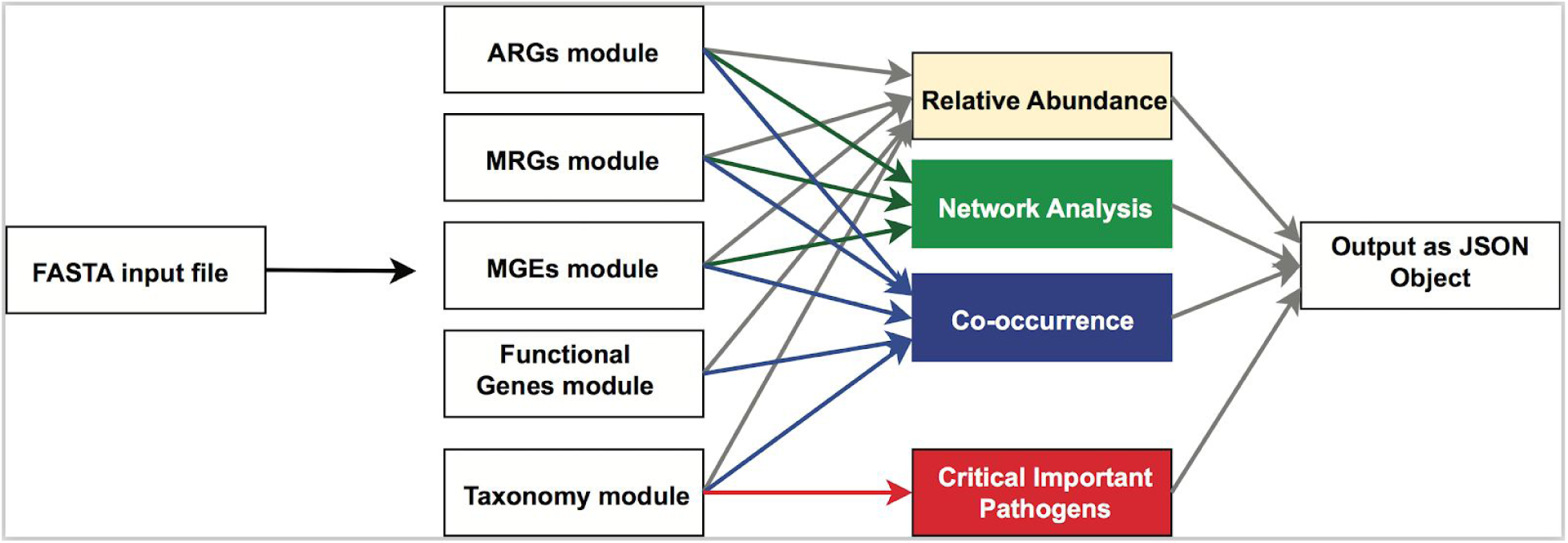
General overview of the NanoARG pipeline. FASTA input reads are processed by five modules to annotate reads according to ARGs, MRGs, MGEs, other functional genes and taxonomy. Annotations are then processed through several stages to obtain the different analysis (relative abundance, network analysis, co-occurrence, and critical important pathogens). All the analysis are packed into a JavaScript Object Notation (JSON) file that can be easily streamed using a http request.

### Clustering of Local Best Hits for Annotating ARGs, MRGs, and MGEs

Traditionally, the analysis of long sequence reads, such as assembled contigs is commonly achieved by first identifying open reading frames (ORFs) within the sequences (44-47) and then searching (e.g., BLAST) with the ORFs against a database to perform functional annotation of the sequences. While nanopore sequences could be considered long contigs, the high sequencing error rate limits the detection of ORFs. Therefore, NanoARG first deploys Diamond (48) for database searches (e.g., for MRG annotation, the reference database is an MRG database) using the nanopore sequences, then clusters all the local best hits into regions, and finally determines the annotation of each region using either the best hit approach or the deepARG prediction (17), as shown in **Figure 4**. Specifically DIAMOND (48) is run with permissive parameters (E-value 1e-5, identity 25%, coverage 40%, and --nk 15000) while bedtools (49) is used to cluster the local best hits in each read into regions. The resulting regions/clusters are then annotated for ARGs, MRGs, and MGEs, as detailed below.

**Figure 4:**
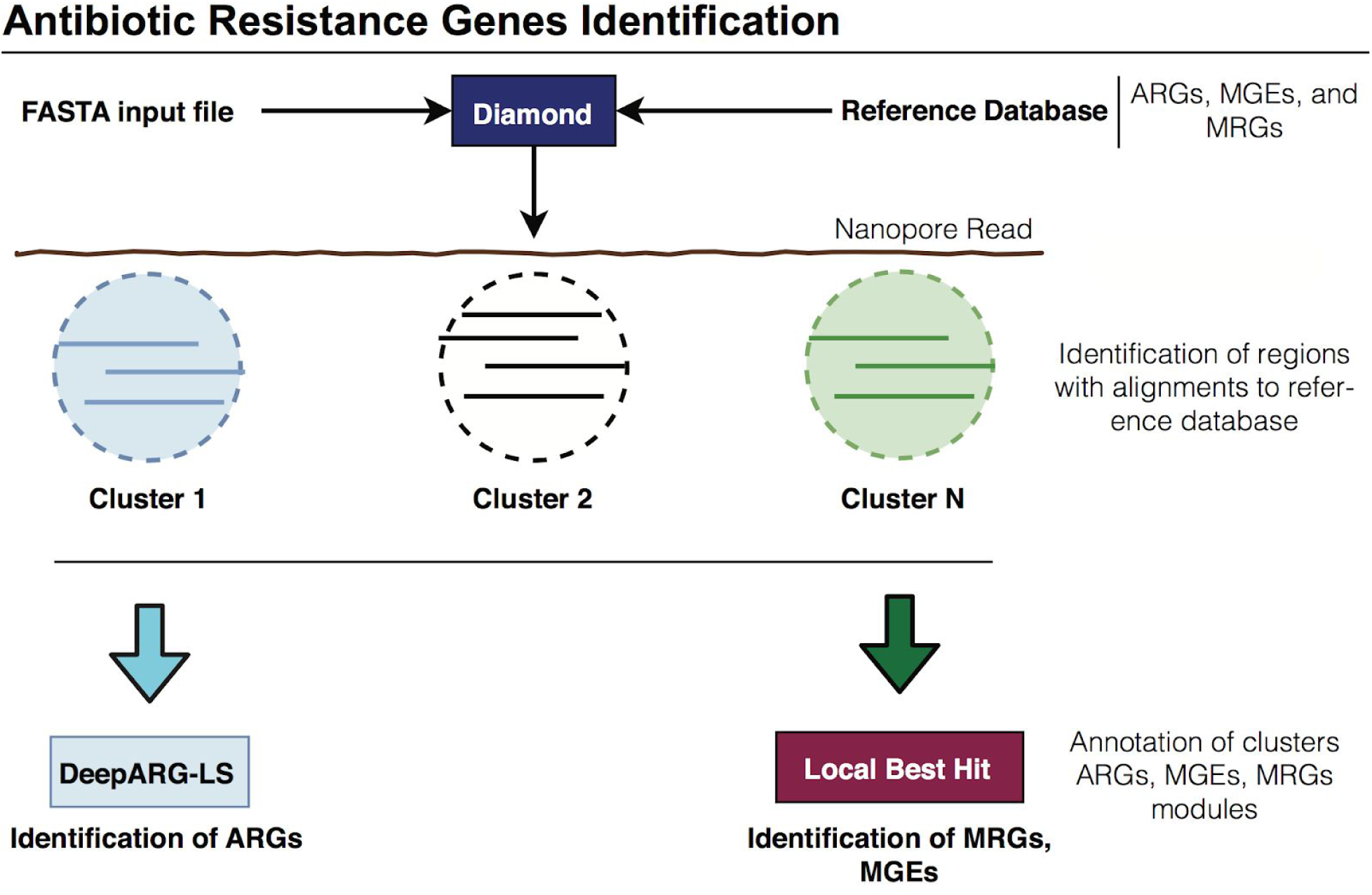
Annotation pipelines. **A)** Identification of ARGs: Input nanopore reads are aligned to the deepARG database using DIAMOND. Alignments are clustered based on their location and annotations are performed using the deepARG-LS model. **B)** Local Best Hit Approach: Identification of the functional genes within the nanopore reads. Alignments are clustered based on their location and the best hit for each cluster is selected, then, alignments are filtered out based on sequence alignment quality.

### ARG Module

Following the clustering procedure of the local best hits to identify putative regions of interest (**Figure 4**), NanoARG uses the deepARG-LS model, a novel deep learning approach developed by Arango-Argoty et al. (17), to detect and quantify ARGs in the regions. A fundamental advantage of the deepARG model is its ability to recognize ARG-like sequences without requiring high sequence identity cutoffs, which is especially useful for nanopore sequences with high sequencing error rates. The deepARG-LS model is applied with permissive parameters, specifically, an identity cutoff of 25%, a coverage of 40%, and a probability of 0.8, to predict that a region corresponds to an ARG.

Abundance of ARG classes and groups is estimated by the copy number of ARGs. To enable the comparison of ARG abundance across samples, analogous to the approach described by Ma et al. (46), the copy number of ARGs is normalized to the total Giga base pairs (Gbp) of the sample to obtain the relative abundance of ARGs:

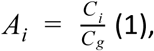

Where *C*_*i*_ corresponds to the total count of ARG *i* (copies of the ARG) and *C*_*g*_ corresponds to the size of the dataset in Gbp, that is, *C*_*g*_ = Γ/μ_*g*_, where Γ is the total number of nucleotides in the library and μ_*g*_ = 1 *x* 10^9^ corresponds to 1 Gbp.

### MRG Module

To annotate MRGs, NanoARG queries the BacMet database (50). Following clustering of the local best hits to identify putative regions of interest (**Figure 4**), NanoARG identifies and categorizes clusters to MRGs according to their best hits. Absolute (copy numbers) and relative abundances of MRGs are computed using **Equation (1)**.

### MGE Database and Annotation Module

Intercellular transfer among bacteria is facilitated via MGEs such as transposons and plasmids (51). Integrons are also key genetic elements of interest as they facilitate capture of multiple ARGS and are often embedded in MGEs, effectively functioning as vehicles for dissemination of multidrug resistance (52). The mechanisms involved in horizontal gene transfer include conjugation, transformation, transduction, and homologous recombination where DNA is incorporated by transposition, replication, and integration (51). Mobile genetic elements were identified from the National Center for Biotechnology Information (NCBI) nonredundant database by using a keyword search (53). Thus, genes related to any of the following keywords — transposase, transposon, integrase, integron, and recombinase— were labeled as associated MGEs. In addition, a set of integrases and class 1 integrons (*Int*I1) were added from the integron-integrase database (54). All sequences were clustered using CD-HIT (55) with an identity of 90%. The resulting MGE database consists of 227,640 genes. Similarly to the annotation strategy adopted for MRGs, nanopore reads are annotated using the MGE database and relative abundance of MGEs is computed using **Equation (1)**.

### Taxonomy Annotation Module

Nanopore reads are classified to their taxonomy lineage using Centrifuge (56), a fast and accurate metagenomics classifier that uses the Burrows-Wheeler transform (BWT) and FM-index. Centrifuge is executed with default parameters (--min-hitlen 25 -f -k 50). Taxonomy relative abundance is estimated in Centrifuge by using an expectation maximization (EM) algorithm similar to the one used in Cufflinks (57) and Sailfish (58). This allows the abundance estimation to be sensitive to genomes that share nearly identical genomic regions. Therefore, each nanopore read is assigned to a particular taxonomy lineage. In addition, nanopore reads not successfully processed by Centrifuge were labeled as unknown.

### Co-occurrence of ARGs, MGEs, and MRGs

The long length nanopore reads offer a unique opportunity to explore the context of ARGs in terms of co-occurrence and potential for mobility. Unlike *de novo* assembly of short reads into longer contigs that might produce chimeric sequences (59), nanopore sequencing yields long sequences naturally thus reducing the potential of chimeras. Therefore, nanopore sequencing is a powerful tool to identify the coexistence of ARGs, MGEs, and MRGs. This analysis is critical to the understanding of antimicrobial dissemination given that co-occurrence and co-selection have been recognized as critically important aspects of ARG spread, where studies have suggested co-occurrence as a mechanism that can facilitate the proliferation of multidrug resistance (60-62). In addition, the co-occurrence of ARGs and MGEs enables tracking of evidence of genetic events of interest, such as HGT (63), along with the identification of novel ARGs, independent from comparison to a reference database (17,64).

To support users in exploring the co-occurrence of ARGs, MGEs, and MRGs in nanopore datasets, NanoARG reports all reads that contain at least one ARG along with its neighboring genes. This data is presented in a tabular format where each entry contains the start position, end position, gene coverage, percent identity, e-value, strand, and taxa corresponding to each read. Furthermore, NanoARG provides a gene map that depicts the gene arrangement, which is useful for visualizing the gene’s co-occurrence and context. Overall co-occurrence patterns are depicted as a network, where nodes represent genes, node sizes represent the number of occurrences, edges between nodes represent genes’ co-occurrence, and edge thickness depicts the number of times the co-occurrence pattern is observed in the data set. Links among nodes are added according to their co-occurrence among the nanopore reads. The network is rendered using cytoscape.js (65).

### World Health Organization Priority Pathogens

The World Health Organization published a list of pathogens that are of particular concern with respect to the spread of antimicrobial resistance (66). This list consists of three priority tiers, namely, critical, high, and medium, as described in **Table 1**. Similarly, the ESKAPE database houses multidrug bacterial pathogens that are critical to human health (67). These two resources are employed by NanoARG to identify the presence of critical pathogens in the nanopore sample. Nanopore reads are matched against the critical pathogens by examining the NCBI Taxa identifier downloaded from the NCBI taxonomy Web site.

**Table 1:** List of twelve families of bacteria considered to be antimicrobial resistant priority pathogens by the World Health Organization (WHO). Pathogens are classified into three categories according to the impact on human health and need for new antibiotics.

| **Importance** | **Pathogen** | **Confer Resistance to** |
| --- | --- | --- |
| **Critical** | Acinetobacter baumannii | carbapenem |
|  | Pseudomonas aeruginosa | carbapenem |
|  | Enterobacteriaceae | carbapenem, ESBL-producing |
| **High** | Enterococcus faecium | vancomycin |
|  | Staphylococcus aureus | methicillin, vancomycin |
|  | Helicobacter pylori | clarithromycin |
|  | Campylobacter spp | fluoroquinolone |
|  | Salmonellae | fluoroquinolone |
|  | Neisseria gonorrhoeae | cephalosporin, fluoroquinolone |
| **Medium** | Streptococcus pneumoniae | penicillin |
|  | Haemophilus influenzae | ampicillin |
|  | Shigella spp | fluoroquinolone |

### Test Run of NanoARG with Eight Nanopore Sequencing Data Sets

To demonstrate NanoARG’s capability for profiling ARGs, four DNA extracts obtained from three different wastewater treatment plant (WWTP) were sequenced using the MinION nanopore sequencing platform and analyzed using NanoARG. In addition, four publicly-available nanopore metagenomic data sets were downloaded and analyzed as detailed below.

### Nanopore Sequencing of WWTP Samples

Four WWTP samples (two influent, two activated sludge) were collected from three WWTPs located in Hong Kong (HK_INF and HK_AS), Switzerland (CHE_INF), and India (IND_AS). Immediately after collection at the WWTPs, samples were kept on ice and transported to the laboratory for processing within 12 hours. Influent samples (1 L) were split into 3 equal volumes, each filtered onto one 0.22 um membrane filter (Millipore, mixed cellulose ester), which was preserved in 1.5 ml 50% ethanol and stored in −20C until DNA extraction. Activated sludge samples (triplicates of 0.5 ml) were mixed with equal volume of 100% ethanol and stored at −20C until DNA extraction. For DNA extraction, filters were torn into small pieces and biomass preserved in ethanol was pelleted by centrifugation (5,000 × g, 10 min) followed removal of ethanol by decanting and pipetting. DNA was extracted with Fast DNA SPIN Kit for Soil (MP Biomedicals) following the manufacturer’s protocol. DNA concentration was quantified with the Qubit dsDNA HS Assay Kit (Thermo Fisher Scientific). DNA for each sample was pooled from triplicate extractions with equal mass. Pooled DNA was purified with Genomic DNA Clean & Concentrator kit (Zymo Research, Irvine, CA). The purity of DNA was then checked using a NanoPhotometer Pearl (Implen, Westlake Village, CA) via the two ratios of A260/280 and A230/260. Each DNA sample (1000 ng) was prepared individually for sequencing using the 1D Native Barcoding Genomic DNA kit (with EXP NBD103 & SQK-LSK108) (Oxford Nanopore Technology) following the manufacturer’s protocol. Each sample was sequenced with a R9.4 flow cell for 24-48 hours without local base calling.

### Heavily Infected Urine Samples (HIU)

Publicly-available MinION data sets obtained from heavily infected urine downloaded from the European Nucleotide Archive (ENA) with the accession study number <u>PRJEB16761</u> (34). Briefly, this library consists of urine obtained from a healthy subject spiked with MDR *E. coli* and an *E. coli* strain cultivated from a heavily infected urine from the same study. The sample was sequenced within 48 hours. Details about the sampling methodology can be found in (34).

### Arctic Glacier Extreme Metagenome (GEM)

This data set was derived from a mixture of samples collected from Arctic cryoconite, located in an unnamed northern outlet glacier in central Svalbard (32). Nine samples were downloaded from the NCBI portal under the accession number PRJEB24565 and were combined into one large file that was further processed on the NanoARG Web service.

### Metagenomic Hospital Fecal Sample (HFS)

This sample consists of DNA extracted from fecal samples of a patient treated with cephalosporins, flucloxacillin, bramycin (an aminoglycoside antibiotic), and colistin (a polymyxin antibiotic) during an ICU care at the University Medical Center Utrecht in the Netherlands. Such treatment was carried out to treat the patient against gut colonization by nosocomial pathogens. In addition, the construction of the metagenomic library was done by a functional metagenomics approach that included a plasmid expression library supplemented with antibiotics, amplified with PCR, and sequenced with the MinION nanopore sequencer. Details about the sequencing protocol can be found in (68).

### Metagenomic Lettuce Spiked *Salmonella* Sample (LSS)

This library was constructed by inoculating *Salmonella* into food samples, including raw chicken breast, iceberg lettuce, black peppercorns, and peanut butter. The MinION nanopore sequencer was used to detect the presence of pathogens and was run for 1.5 hours after enrichment. Details about the library construction can be found at (69).

## RESULTS AND DISCUSSION

NanoARG is an online computational resource designed to process long DNA sequences for the purposes of annotating ARGs, MGEs, MRGs, and taxonomy. A variety of publication-ready figures and tables derived from these annotations can be readily produced facilitating various dimensions of sample comparison.

### Visualization and Data Download

The NanoARG service provides a range of visualization options; including bar charts (**Figure 5A**), tables (**Figure 5B**), gene mapping charts (**Figure 5C**), and co-occurrence networks (**Figure 5D**), displaying individual and combined analyses of ARGs, MGEs, and MRGs. Results can be easily downloaded from the tables and can be configured to include all data, without any filtering. This enables users to deploy their own filtering criteria and customize their analyses.

**Figure 5:**
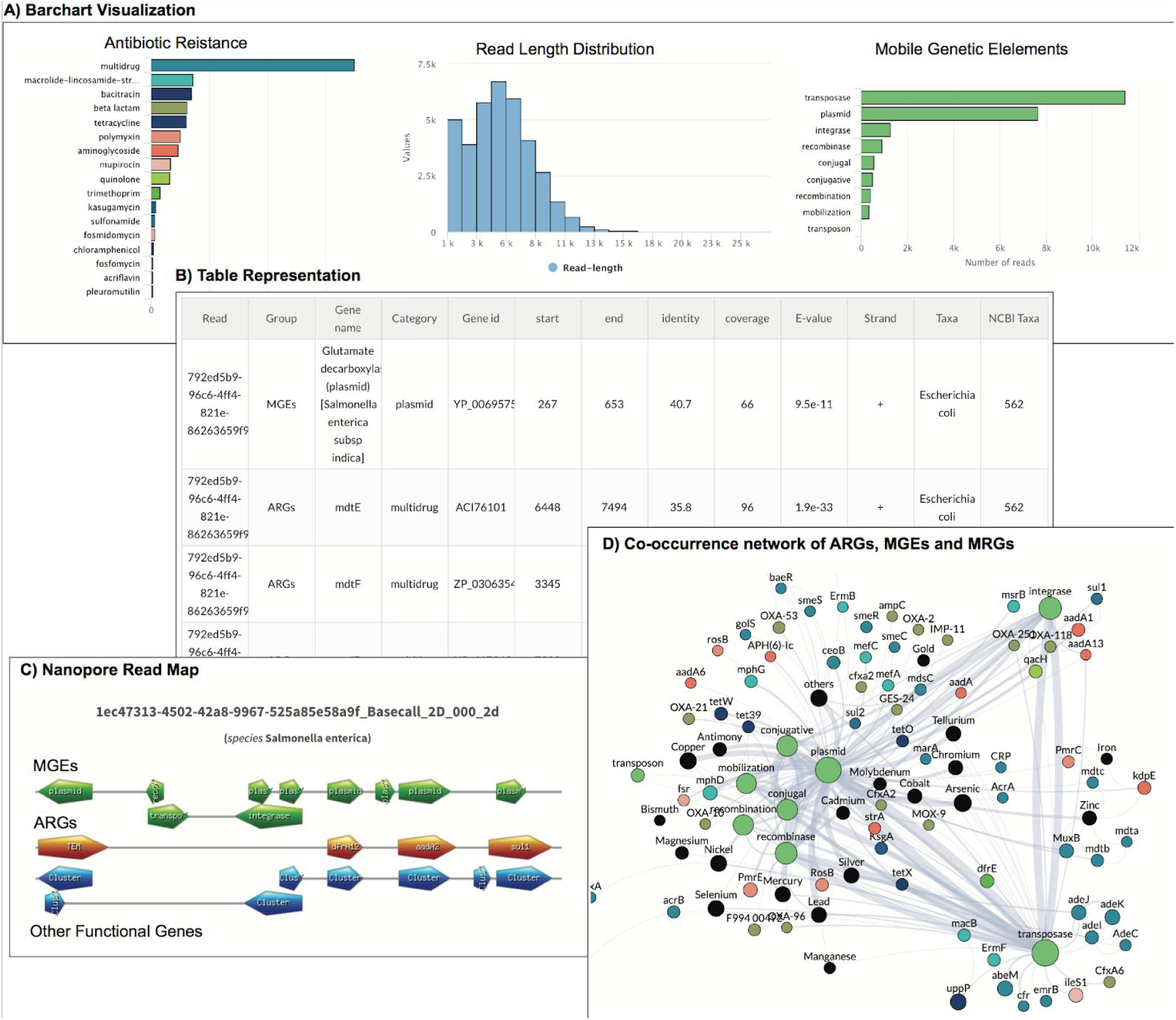
Visualization of NanoARG report. **A)** Absolute abundances (read counts) are shown as barcharts as well as read length distribution and taxonomy counts. **B)** Tabular data: Results are also shown in tables containing all the relevant information for each annotation (E-value, coverage, identity, strand, taxonomy, group, etc.). **C)** Nanopore Read Map: This visualization organize the gene matches in a linear format showing the co-occurrence patterns for each nanopore read with at least one ARG. **D)** Co-occurrence Network of ARGs, MGEs, MRGs: This interactive visualization allows users to drag and drop nodes to visualize the co-occurrence patterns in the sample.

### Effect of Error Correction in the Detection of ARGs

To examine the effect of error correction in the detection of ARGs by nanoARG, nanopore sequences of the HFS sample with and without error correction were used. The complete dataset (Library B) was downloaded from the poreFUME repository including the raw nanopore reads (HFS-raw) along with the corrected reads after the poreFUME pipeline (HFS-poreFUME). In addition, the raw nanopore reads were also corrected (HFS-CANU) using the correction module from the CANU assembler. These three datasets were submitted to the nanoARG pipeline for annotation. Because the HFS dataset does not correspond to a direct metagenomics sample, the ARGs diversity is limited to the ARGs expected by the experimental design. ARGs with high number of hits are referred as “high coverage” ARGs whereas those with few hits are referred as “low coverage” ARGs.

**Figure 6A** shows that the alignment bitscore of all the ARGs is increased after read correction by both canu and poreFUME algorithms compared to the raw uncorrected reads. In particular, for the CANU-correct algorithm, the bitscores of “high coverage” ARGs such as CTX-M, TEM, *aad*A, aac(6’)-I and *erm*B ARGs were significantly improved (**Figure 6B-D**) compared to the raw reads. Similarly, the bitscores of “low coverage” ARGs such as CARB, *erm*F, *fos*A3, *mel*, and *tet*Q also show an improvement after read correction (**Figure 6E-G**).

**Figure 6:**
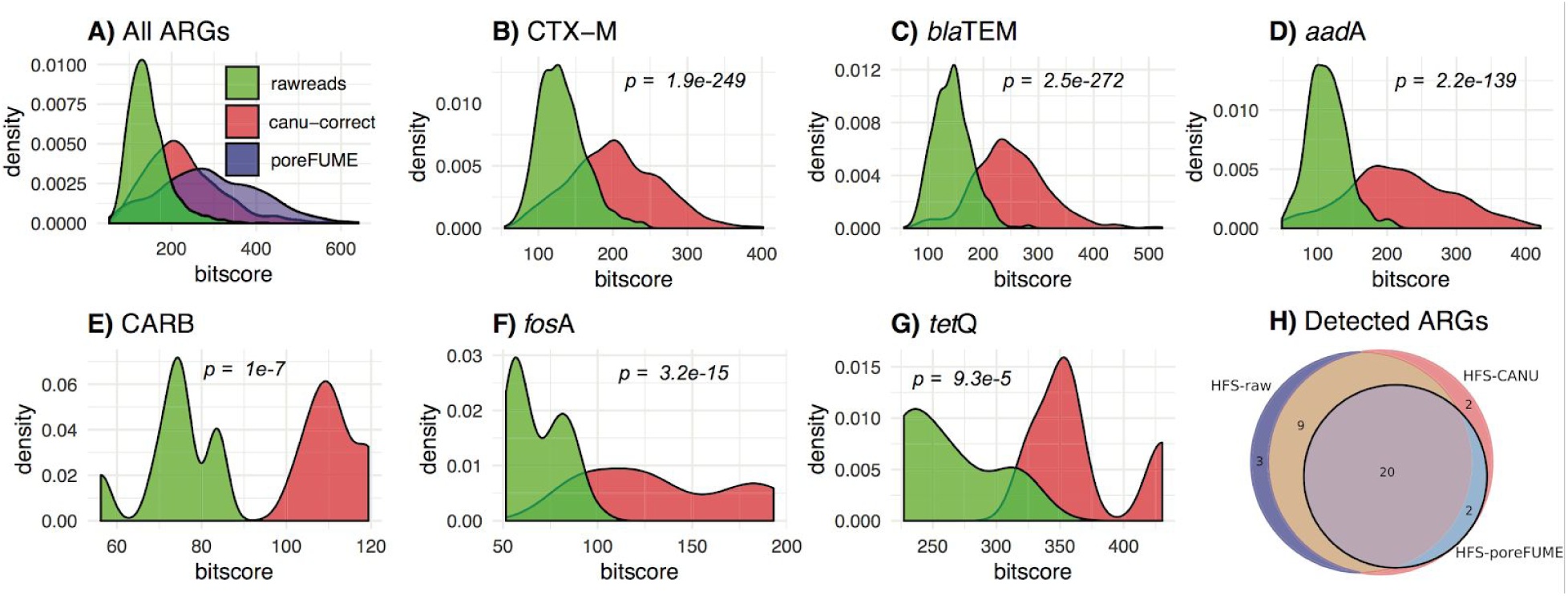
Comparison of error correction in a functional metagenomics sample. Comparison against raw reads and error corrected reads using CANU correct and poreFUME. P-values are computed between the different distributions using a simple T-Test. **A)** bitscore distribution of all ARG alignments. **B-D**) Comparison between raw and corrected reads using CANU correct for ARGs with high depth. **E-F)** Bitscore distribution for raw and corrected reads for low depth ARGs. **G)** Venn diagram showing discovered ARGs by raw and corrected reads by CANU and poreFUME. *Because poreFUME couldn’t ran due to library dependency errors, **Figure 6B-G** contain the transition of quality distribution when comparing CANU-correct and the raw reads

**Figure 6H** depicts the intersection of ARG annotation by nanoARG among the three datasets (HFS-raw, HFS-CANU, HFS-poreFUME). ARGs with a minimum coverage of 80% and an identity greater than 30% were used for this comparison. Altogether 22 unique ARGs were detected in the HFS-poreFUME dataset, 32 for the HFS-raw dataset, and 33 for the HFS-CANU dataset. Out of the 22 ARGs detected in HFS-poreFUME two ARGs (*abe*S and CARB) were not identified in the HFS-raw sample. Further examination revealed that these genes were actually detected in the HFS-raw dataset but were removed after applying the filtering criteria described above. These two genes were also detected after the error correction step (HFS-CANU), indeed, all ARGs that were detected in HSF-poreFUME were also identified after applying the error correction algorithm with CANU. Although there were three uniquely identified ARGs in the HFS-raw dataset (FosC2, LuxR, *emr*K) and four uniquely identified ARGs after CANU correct (CARB, OXY, *abe*S, *van*H), the results show that there was a transition in the annotation from raw to corrected reads. Thus, after comparing the ARG-containing regions within the sequences in the reads, it was observed that those regions were reassigned to ARGs with higher alignment/classification scores after correction. For instance, the raw reads containing the CTX-M gene were reassigned to the OXY gene with higher alignment scores in the HFS-CANU dataset. The CARB gene was detected in both HFS-raw and HFS-CANU datasets. However, the coverage of this gene in the HFS-raw dataset was below the 80% cutoff used for the analysis and therefore removed from the list, whereas it was successfully detected in the HFS-CANU dataset, showing an improvement in the coverage of the alignments. The reads containing the *fos*C2 gene in the HFS-raw sample were reassigned to the *fos*A gene in the HFS-CANU dataset with higher alignment bitscores (73 to 126.3, respectively). Interestingly, the vanH gene was detected exclusively on the HFS-CANU dataset. These results show that the correction step enhances the detection of ARGs in MinION nanopore sequencing samples.

Note that HFS is not a metagenomics sample sequenced directly from the fecal sample. Therefore, to analyze the effect of read correction for metagenomic samples obtained directly from the targeted environment, one WWTP sample was used to analyze the effect of the error correction algorithm. The influent sample from Switzerland (CHE_INF) was processed using CANU correct and submitted along with the raw datasets to nanoARG for annotation. Because of dependency errors present during the execution of the poreFUME pipeline, poreFUME was not performed for the analysis. **Figure 7A** shows the bitscore distribution of the ARG alignments for both raw and corrected reads. It is noticeable that the correction algorithm did not improve significantly (*p*=0.22) the overall ARGs bitscore of the alignments in the WWTP sample. **Figure 7B** shows the intersection of the detected ARGs for the WWTP sample with and without correction, indeed most of ARGs were detected by nanoARG in both raw and corrected reads, only three ARGs were not detected in the raw reads but were detected after read correction (OKP-A, *bcr*A, *otr*C). To see the effect of coverage depth for each ARG, a closer look at the individual ARGs does not show an enhancement of alignment scores for genes with the highest number of hits such as *omp*R, *mex*T (**Figure 7C-D**), as well for ARGs with low number of hits, such as *sul*1, *kdp*E (**Figure 7E-F)**. Because the overlap between the detected ARGs in the raw and corrected reads is greater than 95% (**Figure 7B**), nanoARG does not perform error correction and let users decide whether to upload raw, corrected reads or assembled contigs. In the nanoARG website users can find information on how to perform the error correction using CANU.

**Figure 7:**
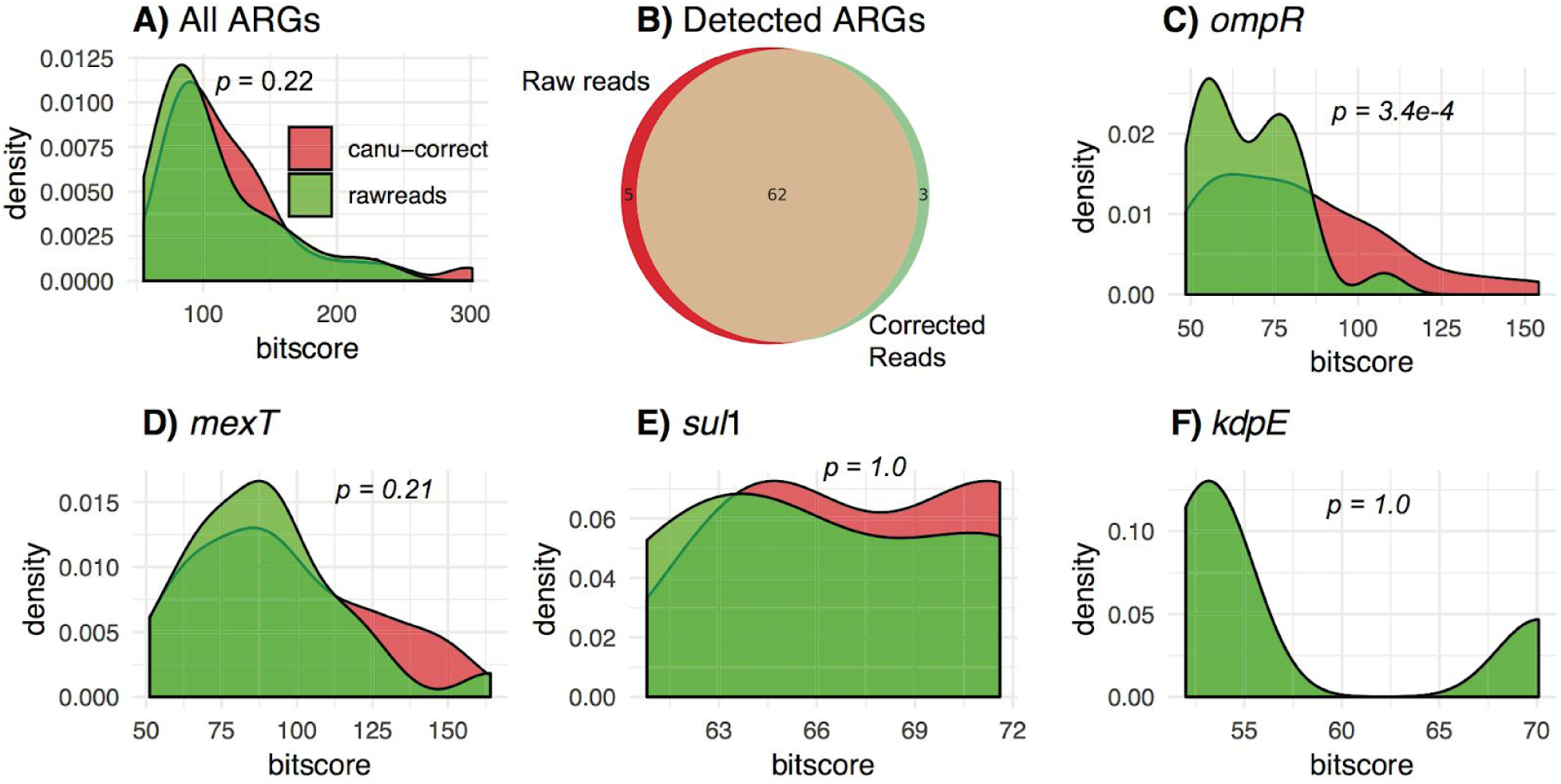
Effect of error correction in an environmental influent WWTP sample. **A)** Bitscore distribution for all ARGs detected by nanoARG using the raw and CANU corrected reads. **B)** Venn diagram showing the intersection of detected ARGs from raw and corrected reads. **C-D)** Examples of the effect of correction in individual ARGs with high number of hits comparing the raw and corrected reads. **E-F)** Effect of correction in ARGs with few hits from the raw and corrected datasets.

To check the effect of time and the consistency for the discovery of ARGs in nanopore samples using nanoARG, several datasets from the LSS sample were analyzed. In detail, the lettuce spiked with *Salmonella* enterica (LSS) study comprised the following datasets: LSS-WGS (whole genome sequencing), LSS-M (shotgun metagenomics), LSS-1.5hN (nanopore sequencing after 1.5 hours) and LSS-48hN (nanopore sequencing after 48 hours). To compare and analyze these datasets, the short reads from LSS_WGS and LSS-M were first assembled using spades with default parameters. Then, the assembled scaffolds were submitted to nanoARG for annotation. On the other hand, MinION nanopore sequencing libraries were first error corrected using CANU correct algorithm and then submitted to the nanoARG pipeline. To evaluate the ability of nanoARG to detect “true” ARGs, nanoARG results from Illumina were compared against the results from the nanopore sequencing. True ARGs are defined here as alignments with an identity greater than 80% and an alignment coverage greater than 90% from the LSS-WGS sample. Thus, a total 28 ARGs pass the filtering criteria. Out of the 28 ARGs, two genes (*mdt*B and *bcr*) were not detected by the shotgun metagenomics illumina sample (LSS-M). When comparing the true ARGs set against the 1.5h nanopore LSS-1.5hN sample, only four true ARGs were detected (aac(6’)-I, *mdf*A, *mdt*G, *mdt*M). This suggests that although nanopore sequencing offers a real time streaming alternative, the detection of specific ARGs would still take several hours. Then, when comparing the true ARGs against the 48h nanopore sample (LSS-15hN), 25 out of the 28 true ARGs were discovered indicating the success of nanopore sequencing for the detection of ARGs. Interestingly, *mdt*B, one of the three undiscovered true ARGs (*mdt*A, *mdt*B and *mdt*C) from the LSS-48hN was not found by either the illumina shotgun metagenomics sample (LSS-M) and the nanopore samples. These three ARGs belong to the same antibiotic resistance mechanism. This analysis shows consistency between the illumina and nanopore sequencing libraries in the detection of ARGs.

### Multiple Sample Comparison: Test Cases

NanoARG provides users with a master table that contains the absolute and relative abundances of ARGs, MRGs, MGEs, and taxonomy annotations for each sample under a particular project. Relative abundances are computed as described in Equation 1. This table includes

### ARG Abundance

WWTP samples contained the greatest number of reads (> 687,835), whereas human-derived samples (HIU, HFS) were comprised of much fewer reads (<67,658) (See **Table 2** for details). Relative abundances of ARGs across the analyzed data sets are compared in **Figure 8**. Notably, human derived samples (HIU, HFS) ranked the greatest number of ARGs per Gb. Particularly, HFS had the highest relative ARG abundance, which is likely related to the sample preparation approach, which intentionally targeted genomic content associated with antibiotic resistance (68). On the other hand, the environmental metagenomic samples ranked much lower ARG relative abundance compared to the hospital and spiked food sample (LSS). Among the WWTP samples, Hong Kong Influent and Hong Kong Effluent ranked the greatest in terms of relative abundance of ARGs.

**Table 2:** Sample collection, metadata and total number of reads for all samples.

| **Sample Name** | **Biome** | **Acronyms** | **Number of Reads** | **Reference** |
| --- | --- | --- | --- | --- |
| **Hong Kong Activated Sludge** | Wastewater | HK_AS | 3307368 | This study |
| **Hong Kong Influent** | Wastewater | HK_INF | 2724813 | This study |
| **Switzerland Influent** | Wastewater | CHE_INF | 687835 | This study |
| **India Activated Sludge** | Wastewater | IND_INF | 1925639 | This study |
| **Arctic Glacier Extreme Metagenome** | Glacier | GEM | 344966 | Edwards, 2016 |
| **Heavily infected Urine** | Human associated | HIU | 36510 | Schmidt, 2017 |
| **Hospital Fecal Sample** | Human associated | HFS | 67658 | van der Helm, 2017 |
| **Lettuce Spiked Salmonella** | Plant surface | LSS | 211806 | Hyeon, 2018 |

**Figure 8:**
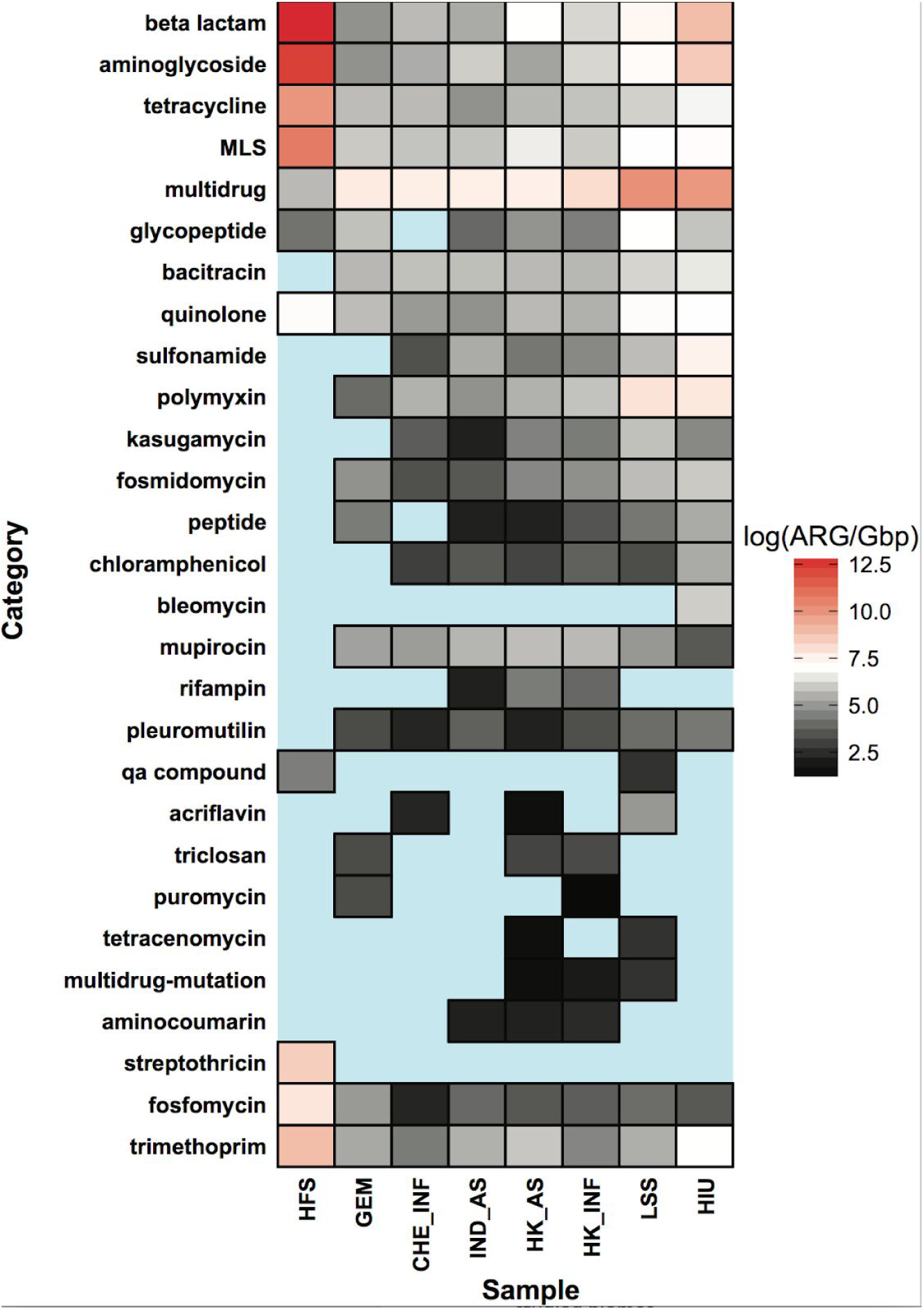
Total relative abundance of antimicrobial resistance genes from the four analyzed biomes. Wastewater treatment plant samples are zoomed to better discriminate differences from the ARG content in each sample.

In considering specific subcategories of resistance, the human fecal sample (HFS) contained the greatest relative abundances of beta lactamase, aminoglycoside, tetracycline, trimethoprim, fosfomycin, streptothricin, quinolone, and MLS antibiotic classes (**Figure 8**). Note that these categories were also prominent in the WWTP and glacier samples, but to a lesser extent than in the hospital derived samples (HIU) and the *Salmonella*-spiked lettuce sample (LSS). In addition, although the multidrug category is highly abundant in HIU and LSS, it was the lowest relative abundance in the HFS sample. Interestingly, although HFS contained the highest relative abundance of total ARGs, the WWTP samples indicated the highest diversity of antibiotic classes compared to other samples. For instance, *sul1* was one of the most prevalent genes in WWTP samples (70), but was not found in the glacier sample. This is consistent with the *sul*1 gene being an anthropogenic marker of antibiotic resistance (71,72). Similarly, the low diversity of beta lactamase genes in GEM (4 beta lactamase ARgs) relative to the WWTP environments (from 25 to 237 beta lactamase ARGs). ARGs from acriflavine, triclosan, aminocoumarin, tetracenomycin, rifampin, and puromycin antibiotic classes were only detected in the WWTP and LSS samples. Interestingly, ARGs derived from acriflavine (a topical antiseptic and disinfectant chemical) were detected in the WWTP activated sludge from Hong Kong, but not the influent. This class is also detected in the influent sample from the Switzerland WWTP. In addition, for the LSS sample, acriflavine ARGs ranked the highest relative abundance. Hong Kong WWTP influent and activated sludge indicated the highest relative abundance of ARGs compared to Indian activated sludge (IND_AS) and Switzerland influent (CHE_INF) (**Figure 9A**). Particularly, the Hong Kong sample showed a decrease in the abundance of multidrug and aminoglycoside resistance genes in the activated sludge sample, but an increase in the beta-lactamase, *MLS*, and trimethoprim ARGs. In addition Indian activated sludge (IND_AS) ranked the highest in relative abundance of sulfonamide ARGs.

**Figure 9:**
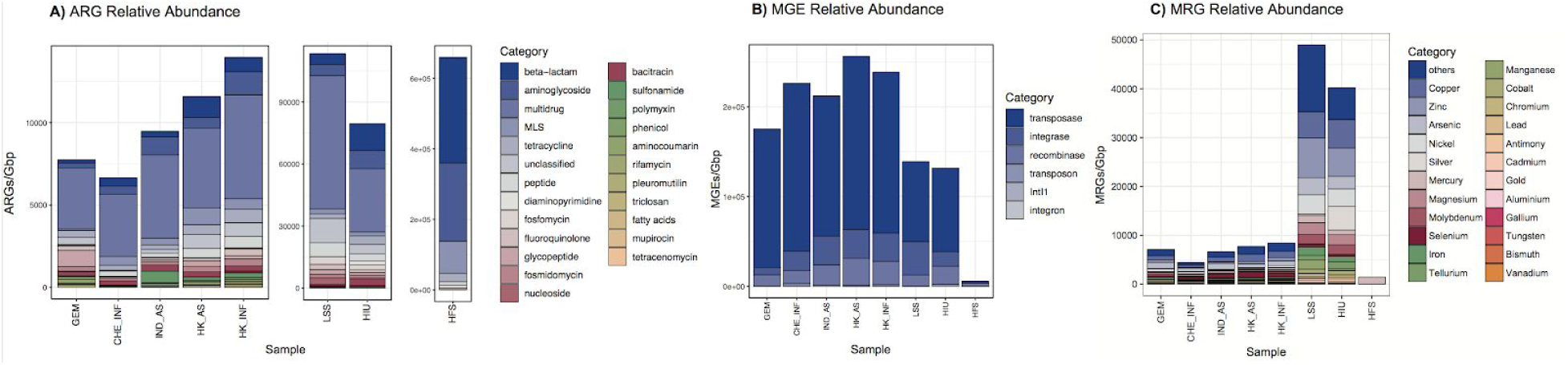
Relative abundance of antibiotic classes for all biomes. Each point corresponds to a particular antibiotic, biome pair. Size and color represent the copy number of ARG divided by 1 Gbp in a logarithmic scale.

### MGE Abundance

NanoARG comprises a collection of genes related to mobility including: transposases, integrases, recombinases, integrons, in addition to a curated database for the class 1 integron *intI*1 (54). Transposases are the prominent MGEs across all samples (**Figure 9B**). Interestingly, the human-derived functional metagenomic sample (HFS) shows the lowest MGEs relative abundance. The *salmonella*-spiked sample along with the heavily infected urine sample, show a lower MGEs relative abundance compared to the environmental samples (WWTP and glacier). Note that the glacier sample GEM has the lowest MGEs abundance compared to the WWTP samples. Interestingly, GEM also has the lowest diversity of MGEs (integrases, transposases, and other MGEs) when compared to other samples. This indicates that bacteria in this environment are not under the same pressure as the microbes from WWTP. On the other hand, a closer look at the class 1 integron *int*I1, one of the major players in the spread of antibiotic resistance (52), shows a different trending. The integron *intI*1 is present in all samples except in the glacier (GEM), likely because glaciers are not under anthropogenic pressure such as antibiotics usage or wastewater discharges. In addition, *int*I1 is ranked with the highest relative abundance on the heavily infected urine sample (HIU) which is expected given the clinical context of the sample. Also, *int*I1 is highly ranked in the India activated sludge sample compared to any other WWTP sample. This suggest that the India sample is under a higher anthropogenic pressure compared to the others WWTP samples resembling the uncontrolled usage of antibiotics in India (73). Noticeable, in the Hong Kong samples, the *intI*1 gene shows a decrease in relative abundance from influent to activated sludge.

### MRG Abundance

MRG profiles were markedly distinct when comparing trends among samples, relative to ARG profiles. The HFS sample has the lowest number of MRGs, with only *merP* and *merT*, part of the mercury transport mechanism (50) detected (**Figure 9C**). In contrast, LSS and HIU samples carried the highest relative abundance of MRGs. Note that HFS is not a direct metagenomic sample, but instead, the raw fecal DNA sample was processed through a plasmid expression library, a transformation cell library, and antibiotic exposure (68). Therefore, the lack of MRGs in HFS could be the result of library preparation. In addition, the HFS sample carried high beta lactamase, aminoglycoside, tetracycline, and MLS abundance, contrasting with low multidrug relative abundance. WWTP samples show a different trend compared to MGEs and ARGs. Specifically, the Switzerland influent sample (CHE_INF) has the lowest relative abundance of MRGs compared to other WWTP samples. Although, CHE_INF has also the lowest ARG relative abundance, its MRG abundance is below half of any other WWTP sample, suggesting that the Switzerland influent WWTP sample has less exposure to heavy metal compounds.

### Taxonomy Profile

The heavily infected urine sample (HIU) shows *Escherichia coli* as the dominant species, which is expected given that this particular metagenome consists of a spiked MDR E. *coli* strain with urine from a healthy patient (34) (see **Figure 10D**). Similarly, the metagenome from food sample (LSS) consists of a *Salmonella enterica* contaminated sample (69). As a result, *Salmonella enterica* is the most abundant followed closely by a *synthetic construct* that refers to artificial samples containing a mixture of different strains (see **Figure 10C**). The results of the HFS sample are interesting because of its library construction. First, the main goal of the study (33) does not consist to build a taxonomy profile, but, to detect ARGs. Thus, the nanopore sequencing is not carried out directly from the source environment, but, it is subject to a specific library preparation (metagenomic sample is transformed into a E. *coli* expression host) that can transform the original microbial composition. The observed taxonomy annotation of HFS consists of a mixture of *E. coli* and other taxas (see **Figure 10B**). Still, NanoARG is able to determine species such as *Escherichia. coli, Klebsiella pneumoniae, Serratia marcescens*, and *Enterococcus faecium*, among others. When performing nanopore sequencing for metagenomes, it is inevitable to observe host DNA. The species distribution in the WWTP samples (see **Figure 10E-H**) shows clearly that these samples contain a large amount of human DNA, in particular, in the Hong Kong WWTP samples, *Homo sapiens* is the dominant species (see **Figure 10F-G**). This host DNA is also observed to a lesser extent in the spiked samples (LSS, HIU). Surprisingly, the HFS sample does not contain any human DNAs, which shows the capabilities of the library preparation, which can be considered as a powerful tool to profile the microbial resistome. Whether it is feasible or not, the use of this strategy for environmental metagenomics needs further examination.

**Figure 10:**
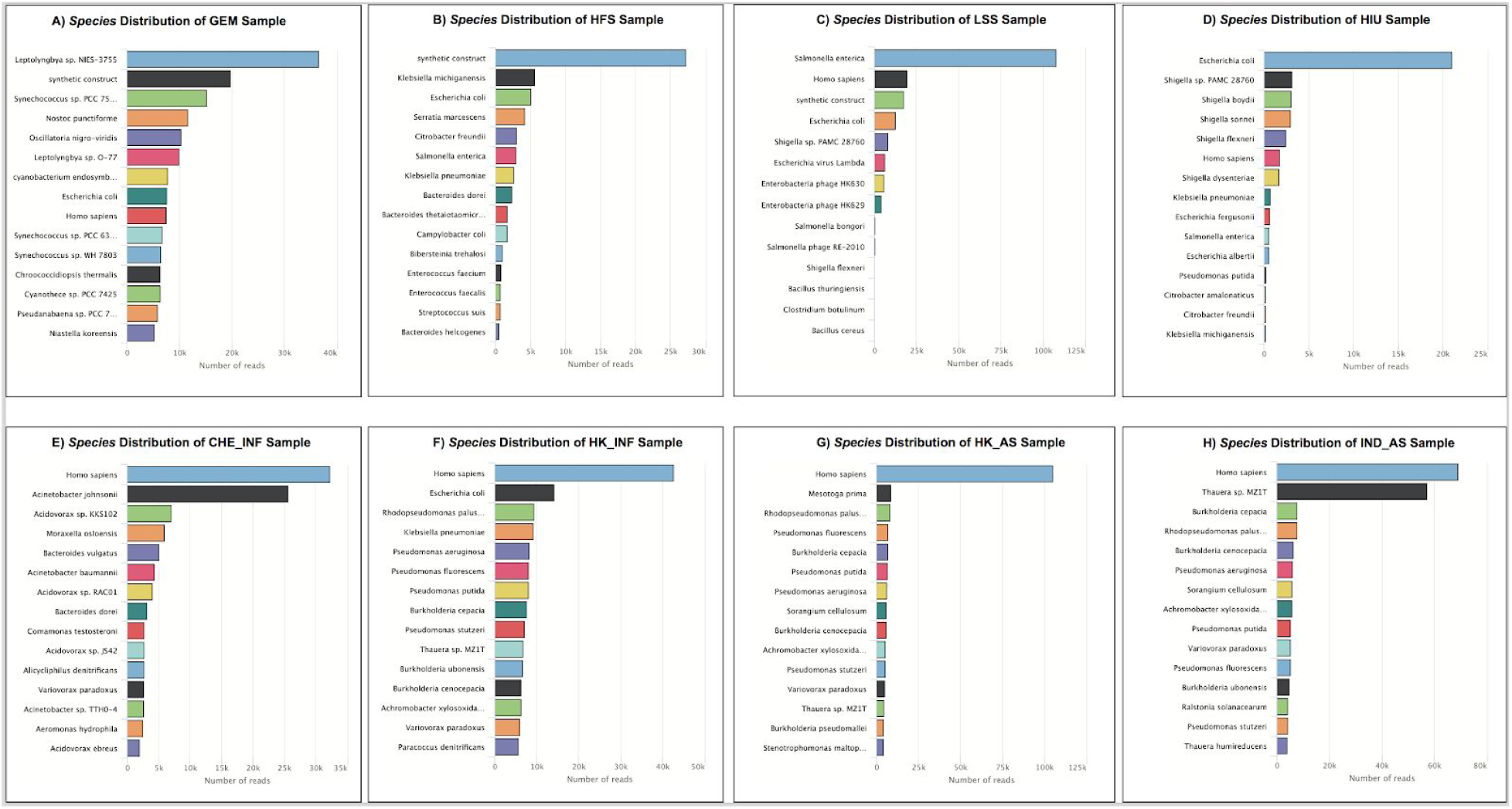
Relative abundance computed as copy of genes by 1Gpb of **A)** Antibiotic resistance classes, **B)** Mobile genetic elements, **C)** Metal resistance genes.

### ARG Context of Nanopore Reads

Long nanopore sequences allow the inspection of ARG linkage patterns and their neighboring contexts. For instance, **Figure 11** shows that the sulfonamide resistance gene *sul1* appears in different contexts depending on the wastewater treatment samples and its host. Also *sul1* is almost all the time around an *integrase/recombinase* along with genes that have been found in plasmids, indicating its great potential in horizontal gene transfer. A commonly observed *sul1* pattern (74), also identified by NanoARG, consists of an *integrase/recombinase* gene, followed by an aminoglycoside (*aadA*) gene, a determinant of quaternary ammonium compound resistance gene (*qacE*), and *sul1*. Interestingly, this pattern seems to be modified in E. *coli* from India and Hong Kong activated sludge environments where the *integrase/recombinase* and the *aadA* region is interrupted by the insertion of a beta lactamase (*OXA*) gene. This linkage pattern differs from the one observed in *Hydrogenophaga sp. PBC* from the Switzerland influent. The contexts of *sul1* in Hong Kong influent and activated sludge samples suggest that the activated sludge process induces the integration of beta lactamase (*OXA*) and aminoglycoside (*aadA, ant3ia*) genes.

**Figure 11:**
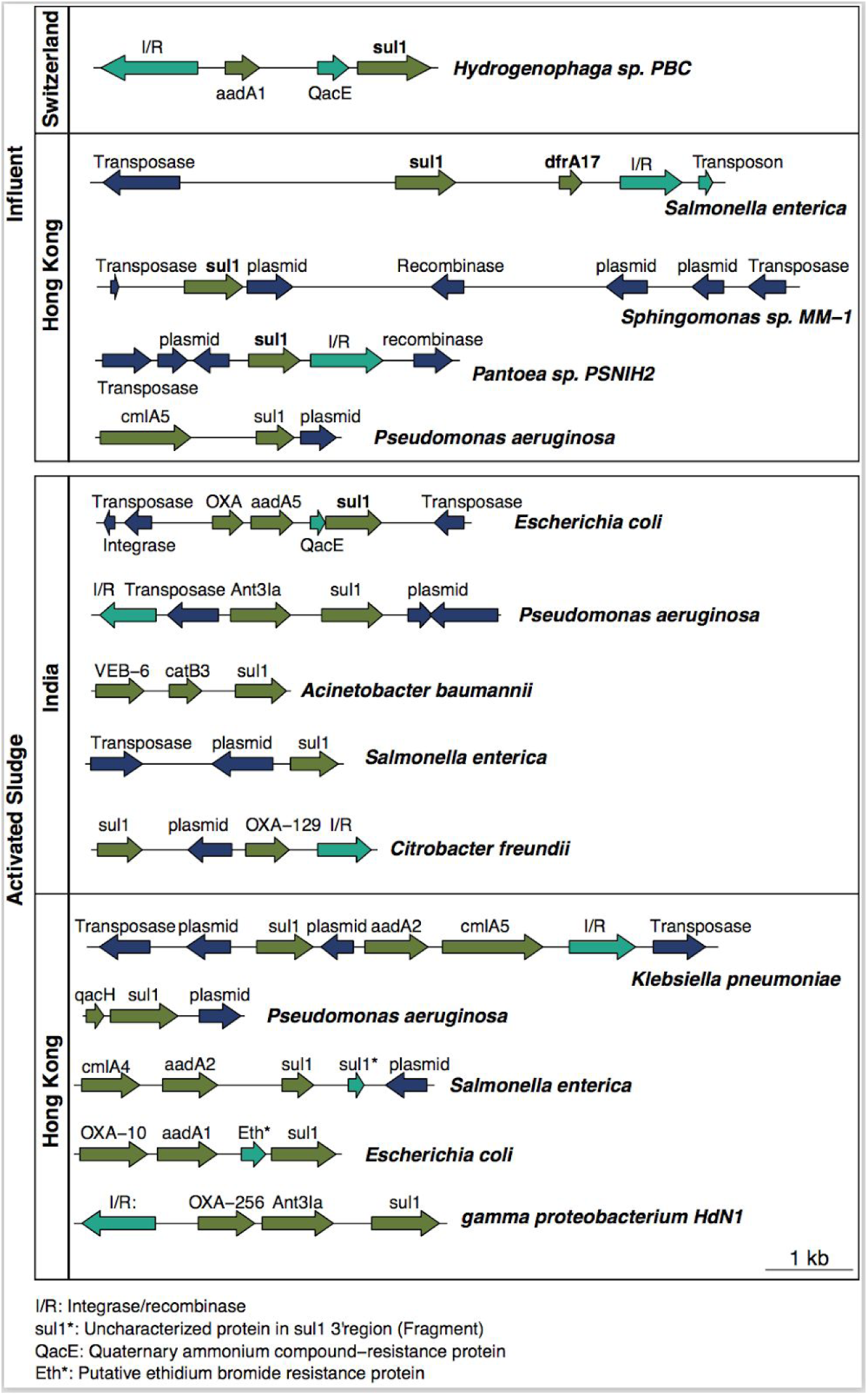
Taxonomy distribution of studied biomes. A) Phylum distribution of WWTP samples. B-H) bar plots with the total number of reads classified at the *Species* taxonomy level of all studied biomes.

**Figure 12** shows the ARG co-occurrence network for all samples. ARGs are linked if they co-occur within the same read and ARGs that appear only once are not shown. Compared to other samples, GEM has the smaller number of ARGs, belonging to only multidrug and trimethoprim classes, and also there is no co-occurrence between these antibiotic categories (**Figure 12A**). Note that WWTP samples show a common pattern of co-occurrence between beta-lactamases and aminoglycoside genes, indicating the high potential of these genes to be carried simultaneously. Interestingly, *sul1* seems to be more abundant in the activated sludge (HK_AS, IND_AS) than in the influent samples (HK_INF and CHE_INF). HFS sample shows that trimethoprim and streptothricin genes have preference to co-occur with the *aad*A1 integron aminoglycoside gene. HFS sample is dominated by aminoglycosides and beta lactamase genes, whereas LSS is dominated by multidrug genes and glycopeptide genes.

**Figure 12:**
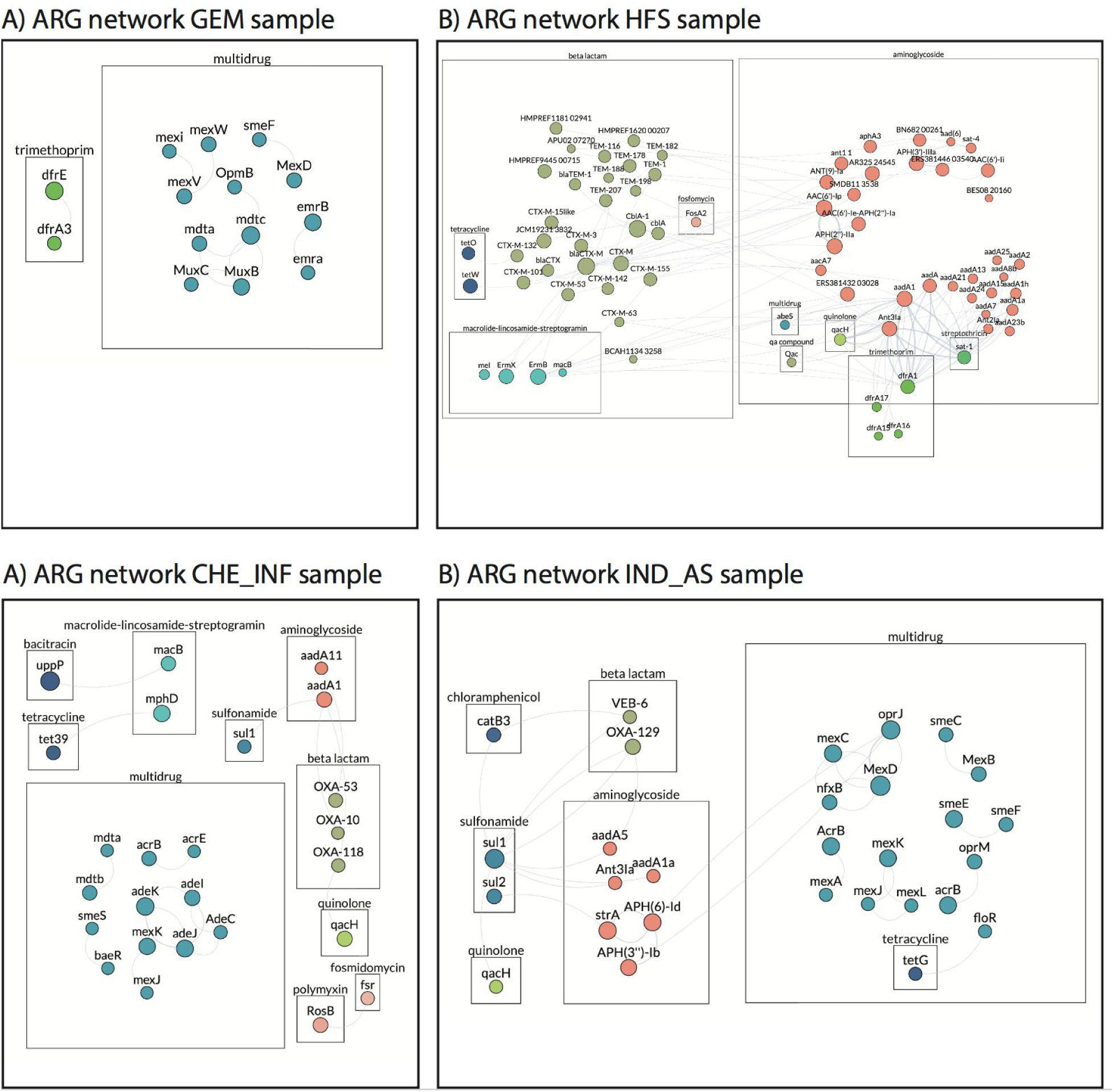
ARG patterns and contexts. Different patterns of ARGs for the wastewater treatment samples (influent and activated sludge). I/R: integrase/recombinase, sul1*: Uncharacterized protein in sul13’ region. *aqcE:* Quaternary ammonium compound-resistance protein, Eth*: Putative ethidium bromide resistance protein.

The *sul1* gene analysis is only one example on how NanoARG facilitates the inspection of any ARG present in the samples. Users can dig deeper different patterns to discover signals of ARG dissemination. The full co-occurrence result can be downloaded for further analysis.

### Critical Bacterial Pathogens

Another important feature of NanoARG is the identification of critical important bacterial pathogens (see **Table 1**) and their association with ARGs. For instance, two of the three most critical pathogens, *Acinetobacter baumannii* and *Pseudomonas aeruginosa*, are found in all wastewater treatment samples (see **Table 3**), whereas *Enterobacteriaceae* (carbapenem-resistant pathogen) is found only in the Hong Kong Influent sample. This pathogen carries a multidrug (*mdtD*), an aminoglycoside (*strA*) and a recombinase gene. In addition, the Hong Kong influent sample contains the *Neisseria gonorrhoeae* pathogen that carries the *mtrE, mtrD, mtrC, cpxr* (multidrug), *pbp2b* (beta lactamase), and uppP (bacitracin) genes. However, no MGEs are found along with these ARGs, indicating no imminent risk of lateral transfer of ARGs. *Pseudomonas aeruginosa* seems to be the most abundant critical pathogen across all samples and is particularly abundant in the India activated sludge sample. Interestingly, no pathogens are found in the glacier (GEM) sample.

**Table 3:** List of critical important bacterial pathogens found in the wastewater treatment samples.

The analysis of critical important pathogens in NanoARG provides a way to narrow down the analysis of ARGs for information considered a top priority for human health. Keeping track of ARGs in pathogens and their associations in different environments can help the understanding of the dissemination of ARGs.

### Data Access and Web Service

As shown above, the NanoARG server provides users with a wide variety of analyses for ARG, MGE, MRG, and taxonomy annotations. NanoARG also provides several types of visualizations to interpret the results. In addition, NanoARG allows users to download the results (e.g., absolute/relative abundances, co-occurrence network associations, taxonomy annotations, and ARG context patterns) in a tabular format containing the fields required for tuning the results (E-value, identity percentage, and coverage). These tables can be used for further processing and statistical analysis. NanoARG core computation is carried out by a high performance computing cluster at Virginia Tech under the Advanced Research Computing center (http://arc.vt.edu), enabling fast and efficient processing of the nanopore reads. The NanoARG Website is developed using the Google Angular 5 framework (https://angular.io), the back-end is developed under the node.js framework (https://nodejs.org/en/). Finally, the computing pipeline is developed using the Luigi framework, allowing the monitoring and rescheduling of jobs failed during execution (https://github.com/spotify/luigi).

### NanoARG Usage Recommendation

Note that the various analyses provided by NanoARG are not restricted to nanopore sequencing reads. In fact, NanoARG can be applied to any set of long sequences. For instance, sequences from different technologies such as PacBio long-read sequencing or assembled contigs from short sequencing reads can be directly processed in NanoARG. Depending on specific research needs, different studies may have different requirements, e.g., some require more stringent criteria whereas others less. Thus, to allow for flexibility and customization, NanoARG provides users results produced by relaxed annotation parameters so that users can filter the results further for their own analyses.

## CONCLUSIONS

NanoARG is a public Web service dedicated to the analysis of antibiotic resistance genes from nanopore MinION metagenomes. This platform has been developed to analyze environmental metagenomes from MinION nanopore sequencing reads. However, it is not restricted to this type of data. As shown in the analysis of real data, NanoARG can be used to profile the antimicrobial resistance of any biome. Indeed, NanoARG could be used to analyze any set of long sequences (e.g., assembled contigs). Its user friendly interface makes it easy to process a new sample as well as retrieve the results. Unlike other services such as WIMP (What Is In My Pocket) dedicated exclusively to antimicrobial resistance, NanoARG offers the analysis of heavy metal resistance genes, mobile genetic elements, taxonomy annotation, identification of critical pathogens, and co-occurrence networks. In addition, NanoARG is the first Web service devoted to the analysis of environmental samples. This pipeline uses deepARG, a novel approach for identifying ARGs from metagenomes and implements a local strategy to annotate genes from long nanopore reads. It uses a set of permissive parameters that allows high flexibility to detect homologous genes, which is important because of the high error rate within nanopore sequences.

## Supporting information

